# Direct imaging of the circular chromosome in a live bacterium

**DOI:** 10.1101/246389

**Authors:** Fabai Wu, Aleksandre Japaridze, Xuan Zheng, Jacob W. J. Kerssemakers, Cees Dekker

## Abstract

New assays for quantitative imaging^1–6^ and sequencing^7–11^ have yielded great progress towards understanding the organizational principles of chromosomes. Yet, even for the well-studied model bacterium *Escherichia coli*, many basic questions remain unresolved regarding chromosomal (sub-)structure^2,11^, its mechanics^1,2,12^ and dynamics^13,14^, and the link between structure and function^1,15,16^. Here we resolve the spatial organization of the circular chromosome of bacteria by directly imaging the chromosome in live *E. coli* cells with a broadened cell shape. The chromosome was observed to exhibit a torus topology with a 4.2 μm toroidal length and 0.4 μm bundle thickness. On average, the DNA density along the chromosome shows dense right and left arms that branch from a lower-density origin of replication, and are connected at the terminus of replication by an ultrathin flexible string of DNA. At the single-cell level, the DNA density along the torus is found to be strikingly heterogeneous, with blob-like Mbp-size domains that undergo major dynamic rearrangements, splitting and merging at a minute timescale. We show that prominent domain boundaries at the terminus and origin of replication are induced by MatP proteins, while weaker transient domain boundaries are facilitated by the global transcription regulators HU and Fis. These findings provide an architectural basis for the understanding of the spatial organization of bacterial genomes.

---

It is increasingly understood that the spatial organization of a genome is crucially important for its biological function. Recent experiments have led to different proposals for the structure of the highly compacted 4.6-Mbp circular genome of *E. coli* (Fig.1a). For example, while the macrodomain model suggested large (~0.5-1Mbp) domains of internally interacting DNA brought together by site-specific systems, including a condensed-terminal domain^11,17,18^, the linear filament model depicted a rather uniformly stacked nucleoid body connected by a thin terminal string^2,19^. Whole-chromosome imaging^1,20^ would be ideal to resolve and characterize the fine structure of chromosomes. Unfortunately, such studies suffer from a limited resolution because the chromosome is tightly confined within the rod-shape cell that is narrower than 1 μm, and furthermore because the chromosome continuously replicates upon cell growth. We set out to overcome these limitations by, respectively, (i) using a drug (A22, an MreB-inhibitor) which widens the cell while keeping it alive in a fully physiologically active state^21^, and (ii) using a strain (with a *dnaC2(ts)* allel^22^) that stops initiating DNA replication at 40°C (Methods). As a result, cells maintain only one single chromosome while growing from a rod into a lemon shape (~2-μm wide, ~4μm long, and ~1-μm high under an agarose pad) over the course of 2-3 hours (Fig.1b).

**Figure 1.**
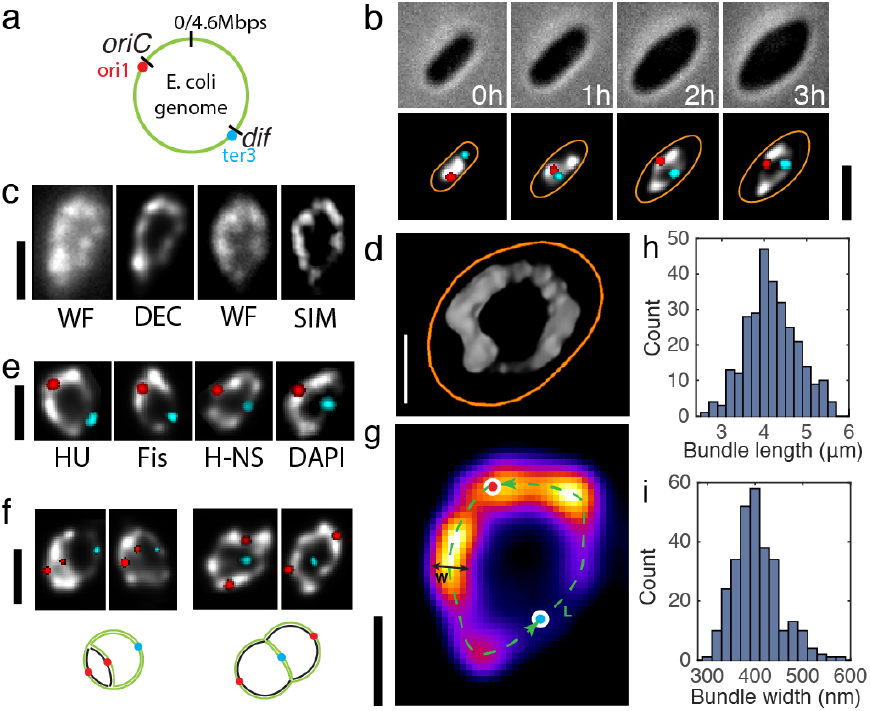
**The circular *E. coli* chromosome exhibits a toroidal donut shape that can be visualized upon cell expansion.** a. Schematic of a *E. coli* genome. Two fiducial markers are shown in red (Ori1 marked by lacO arrays, targed by LacI-mCherry) and cyan (Ter3 marked by tetO arrays, targeted by TetR-mCerulean).
b. Time-lapse fluorescence images of an *E. coli* (*dnaC2(ts)* allel) cell growing into a lemon shape at 40 °C under A22 treatment. Top panel, phase contrast image; bottom panel, overlay of Ori focus (red) and Ter focus (cyan) on a grey-scale 3D-deconvolved image of the chromosome labeled by HU-mYPet. Time is indicated in hours.
c. Fluorescence images showing two opened circular chromosomes captured by different methods. Left: WF, widefield image, and DEC, 3D-deconvolved image of that WF. Right: WF and SIM, structured illumination microscopy image of that WF image.
d. Donut-shape chromosome of E coli, as imaged in 3D-reconstructed SIM. Orange outlines the cell contour.
e. Similar donut-shape chromosome images are obtained for different DNA-binding fluorescent labels (HU-mYPet, Fis-mYpet, H-NS-mYPet, DAPI; all DEC images).
f. Fluorescence images of early (left) and late (right) stages of DNA replication of circular chromosomes. Bottom: cartoon illustrations; black strands indicate newly replicated DNA.
g. Fluorescent image of a donut-shape E. coli genome shown as a heat map. Indicated are the ridge of the bundle (green dashed line), the oriC and dif genomic loci near the origin and terminus of replication (red and blue dots respectively), and the bundle width (black line).
h. Histogram of chromosome bundle lengths measured along the bundle ridge (cf. panel g). n=292.
i. Histogram of the average chromosome bundle widths quantified as the full-width-at-half-maximum of the peak intensity across the donut chromosomes. n=292.Scale bars in b/c/e/f, 2 μm. Scale bars in d/g, 1 μm.

Interestingly, upon a two-fold widening of the cell, the single *E. coli* chromosome was observed to laterally expand and gradually open up into a torus (Fig.1b). This topology was consistently observed through different imaging techniques such as wide-field epifluorescence and Structure Illumination Microscopy (SIM) (Fig.1c-d, Fig.S1-2, Methods), and with different fluorescent labels in live cells (Fig.1e). Note that the toroidal geometry is not trivial, since *a priori* other outcomes could also have been expected for a single chromosome upon cell widening, such as a homogenously spread-out globular cloud of DNA^23^, or a nucleoid that would maintain its shape as in rod-shaped cells^2^. Our images of an open ring-like geometry agree with previous proposals^2,4,9,19^ that the circular *E. coli* chromosome consists of two independent arms flanking the origin of replication, an arrangement distinct from the SMC-mediated arm zipping that was reported for *Bacillus subtilis*^*7*^ and *Caulobacter crecentus*^10^. Interestingly, the chromosomes were able to immediately resume replication and cell division after a brief reactivation of the DnaC proteins (Fig.S3), during which the replicated regions branched out while conserving their bundle morphology (Fig.1f). By contrast, upon treatment with drugs such as rifampicin (which blocks transcription by inhibiting RNA Polymerase) or ciprofloxacin (which impedes the homeostasis of supercoiling through inhibiting gyrase and TopoIV activity), the chromosomes collapsed and generally lost the torus topology (Fig.S3). We conclude that the torus topology is maintained by active physiological processes, and therefore serves as an excellent model object for resolving the organizational principles of a *E. coli* chromosome in live cells.

The direct visualization of the genome allowed us to quantitatively measure the width and length of the *E. coli* chromosome bundle (Fig.1g-i). Facilitated by 3D-deconvolution which reduced the out-of-focus background intensity in wide-field imaging (Fig.1c, Fig.S2), we mapped the ridge line of the chromosome, and measured the length along this contour. The chromosome contour length was found to be 4.2 ± 0.6 μm (mean±s.d., Fig.1h, Fig.S2, n=292, Methods), while the average bundle thickness, characterized by the full-width-at-half-maximum (FWHM), was 0.40 ± 0.05 μm (Fig.1i). The intrinsic 4.2-μm length and 0.40-μm thickness provide a quantitative basis for future modeling of the polymer structure and the mechanics of the chromosome as well as the role of confinement and volume exclusion in the segregation of newly replicated DNA^1,2,16^.

The donut-shape chromosomes were observed to be strikingly nonuniform. The DNA density was not homogenous along the circumference, but, unexpectedly, partitioned into blob-like domain structures (Fig.2a). Using a custom-made blob-analysis script (Methods, Fig.S4), we found that each chromosome contains between 3 and 7 apparent domains, with 4 as the most probable number in a distribution described by a lognormal probability density function (PDF) (Fig.2b). The blobs showed a broad range of physical sizes, with a diameter *D* ranging from 200 nm to 1 μm (mean±s.d. = 0.56±0.18 μm, Fig.2c). Next, we quantified the DNA length *L* (measured in base pairs) contained in each cluster based on fluorescence intensity. This yielded values from 300 kbp to above 3 Mbp (Fig.2d), showing a lognormal PDF with a peak at 1.2 Mbp (Fig. 2d). *D* was found to scale with *L* according to a *D ~ L*^*α*^ power law with an exponent α = 0.39 ± 0.05 (Fig.2e), a scaling property that is surprisingly similar to that of heterochromatin in *Drosophila*^6^.

**Figure 2.**
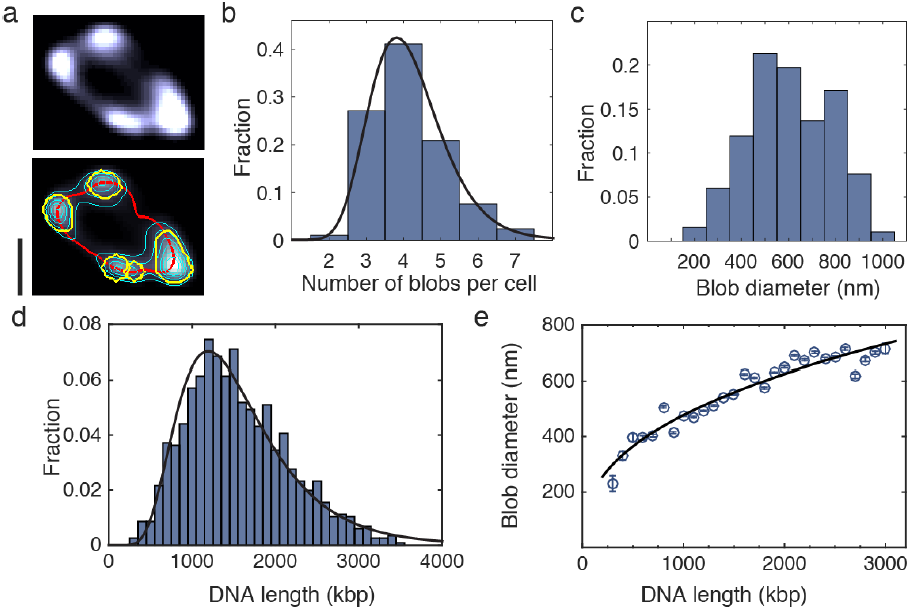
**Domain distribution within the circular chromosome of *E. coli*.** a. Example of the automated domain recognition. Top panel, fluorescent image of a toroidal chromosome. Bottom panel, same image with ridge line (red), equal-intensity lines (thin cyan lines), and blob boundaries (yellow). Scale bar, 2 μm
b. Number of blobs per chromosome. Black line shows the probability density function (PDF) of a fitted lognormal distribution. n=292.
c. Bar plot of the PDF of the physical size (diameter) of the blobs.
d. Distribution of domain sizes (DNA length in kbp) in all measured cells. A fitted lognormal PDF is shown in black.
e. Blob diameter *F* versus DNA length *L* contained in a blob. Blue circles and error bars indicate mean ± s.e.m. calculated for all cells within a 100 kbp bin size. Black line is a fit of a power law, *F ~ L*^*α*^ (α = 0.39 ± 0.05).

In order to quantify the average DNA density as a function of the genomic sequence coordinate, we mapped the fluorescence intensity along the ridges of the donut-shape chromosomes (Methods, Fig. S5), aided by fiducial markers (Fig. 1a and 3a). Figure 3a-c shows the data from a strain with labels at the L3 and R3 positions^19^, which divide the circular chromosome into an *oriC*- and *dif*-containing branch that have an intensity ratio of 72:28% (n=84, standard error 1%), close to the expected 70:30% DNA ratio. Next, we mapped the contour coordinates onto the genome sequence by constructing a cell-average cumulative density function that starts and ends at the *L3* foci, which allowed physical positioning of the genomic loci including *oriC* and *dif* sites (where DNA replication initiates and terminates, respectively) onto the torus (Fig.3d). This yielded the DNA density distribution along the chromosomal torus, averaged over all cells, see Fig.3e. The average DNA density is seen to display a pronounced M-shape curve with a very deep minimum located at the *dif* locus and a second, less deep yet well developed, minimum at the *oriC* locus (Fig.3e). These *oriC* and *dif* loci are connected by the DNA-dense left and right arms, which show a slight asymmetry with a slightly higher DNA density peak in the left arm. The global M-shape as well as the same locations of the minima were also found in a second independent strain with Ori1 and Ter3 labels located adjacent to the *oriC* and *dif* sites, respectively (Fig.1b, 3f, Fig.S6), albeit with less well distinguishable left/right arms due to symmetry (Fig.1b), and were also conserved in all fluorescent labels (Fig.S7).

**Figure 3.**
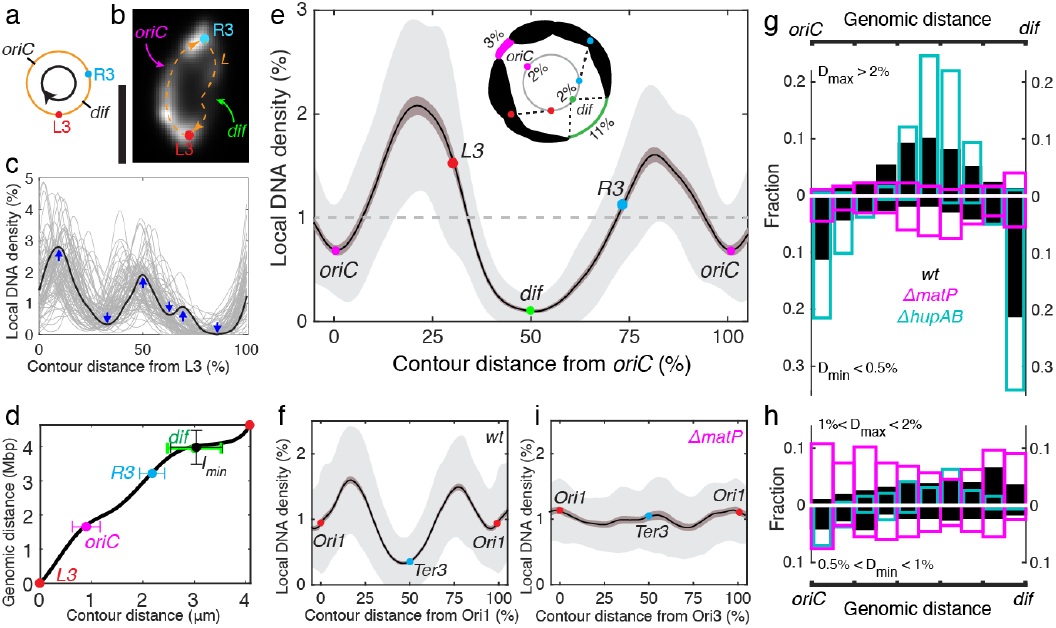
**DNA density mapping along the circular *E. coli* genome.** a. Schematic of the *E. coli* genome with two fiducial markers at the left and right arms shown in b-e. The arrow indicates the direction of density mapping.
b. Example of a circular chromosome as indicated in a, with the HU-mYPet fluorescence intensity shown in grey scale. Scale bar, 2 μm.
c. Local DNA density along the ridge line of single circular chromosomes plotted as a function of percentile distance from L3. Each line indicates a single chromosome. 1 random example is highlighted in black. Blue arrows indicate local maxima (peaks) and local minima (valleys) in this example curve. n=82.
d. Average cumulative density function mapping the genomic coordinates to the contour coordinates along the ridge of the circular chromosome. Marks indicate the measured positions of the R3 locus and global minimum *I*_*min*_, and the predicted positions of *oriC* and *dif* sites. Error bars indicate s.d‥
e. Local DNA density plotted versus genomic coordinate in percentile distance along the ridge line from the predicted *oriC* sites, with mean values and positions of *dif*, *L3* and *R3* indicated. Dark and light shading indicates s.e.m and s.d. Inset: schematic illustrating the DNA density distribution along a typical circular *E. coli* chromosome as concluded from the blob analysis and contour density analysis. n=292.
f. DNA density distribution of a second independent strain with Ori1/Ter3 labels, plotted as in panel e. Note that in this strain left and right arms can be distinguished less well due to symmetry (Fig. 1b). n=74.
g-h. Distributions of local maxima (plotted upwards) and local minima (downwards) in the DNA intensity within individual chromosomes along the genomes from *oriC* to *dif* site. Bin size 5% of the genome length (230kbp). Local maxima and minima are identified as in c. ‘D’ denotes local DNA density.
i. DNA density distribution of a *ΔmatP* strain with Ori1/Ter3 labels, plotted as in panel f.

Individual DNA density plots of single chromosomes (cf. Fig.3c) exhibited a variety of local maxima and minima, from which we extracted the genomic locations of domain centers (high DNA density) and domain boundaries (low DNA density), respectively. As shown in Fig.3g, very pronounced domain boundaries were consistently found at the *oriC*- and *dif*-proximal regions, whereas the prominent domain centers were found near the centers of the left/right arms. By contrast, less pronounced domain centers and boundaries were found to distribute more evenly throughout the genome (Fig.3h). This led us to hypothesize that different mechanisms may be at play in defining the chromosomal domain structure in *E. coli*: (i) a mechanism that reduces DNA condensation at the *oriC* and *dif* regions or promotes interactions at the centers of the two arms, and (ii) a mechanism causing transient dynamic domains across the genome – both of which we will discuss now.

First, we examined the origin of the pronounced gap in the DNA density near the terminus region (Fig.3e-g, Fig.S6). We found that near the *dif* site, where the global density minima consistently reside (Fig.3d), a mere 2% of the genome (92kbp) spans as much as 11±6% of the physical bundle length (Fig.3e, inset). This region posits at the center of the previously proposed 800kbp Ter macrodomain, which was proposed to be condensed by MatP proteins through their binding to the *matS* sites^13,17,24^. We therefore examined the DNA density distribution in Δ*matP* cells, which also opened up into a torus topology. We found that, strikingly, all pronounced peaks and gaps disappeared in the average density distribution (Fig.3i), and peaks and gaps in individual cells were evenly distributed across the chromosomes with no prominent features in either the *oriC* or *dif* sites (Fig.3g-h, Fig.S8). MatP thus is found to be crucial for the formation of the prominent domain boundaries at both the *dif* and *oriC* regions. Interestingly, MatP drives the *dif*-proximal region into a thin string^2,15,19^ rather than condensing it as previously proposed^17,24^, likely for promoting its accessibility to division proteins, topoisomerases, and DNA translocases during terminal segregation^25,26^. Indeed, the thin terminal string persisted at different stages of the replication cycle (Fig.1f) in wildtype, but not in Δ*matP* cells (Fig.S8). The impact of MatP on the *oriC* region is in line with the recent finding that MatP interacts with MukBEF^27^, possibly enabling the accessibility of such factors involved in regulating the initiation of DNA replication and segregation.

Although the average DNA density distribution clearly showed 2 peaks at the two chromosomal arms (Fig.3e), individual chromosomes typically contain a larger number (3-7) of physical domains (Fig.2b). We examined the origin of the secondary domain boundaries by quantifying their distribution in the presence and absence of the nucleoid-associated proteins (NAPs) HU, H-NS, and Fis, which function as global transcription regulators^28^. Whereas all NAP mutants preserved the overall M-shape DNA density distribution with a deep minimum at *dif* region and a less pronounced minimum at *oriC* region (Fig.S7), deletion of NAPs could affect the domain boundaries: While eliminating H-NS proteins had little effect, omitting HU and Fis proteins led to a very significant (near 80%) loss of the domain boundaries in the central regions of the two arms, which instead became significantly more enriched with domain centers (Fig.3g-h, and see Fig.S9 for a detailed analysis). Given that HU and Fis are transcription activators that localize sequence-nonspecifically throughout the genome (Fig.S7, S9), they likely do not produce domain boundaries directly, but instead promote their formation indirectly by stabilizing supercoils^29,30^ within domains. These data, as well as our observation that transcription globally expands chromosomes (Fig.S2), indicate that transcription machineries process DNA to dynamically partition chromosome into domains of various sizes. This view aligns with emerging data from *C. crescentus*^8^ and eukaryotes^5,6^.

Indeed, very pronounced dynamics are apparent in time-lapse imaging of the donut-shape chromosomes. Figure 4a shows an example of a SIM movie that displays remarkable up-to-Mbp re-arrangements in the chromosome morphology at a sub-minute time scale (see Video 1). Our approach allowed us to construct proximity maps of a single genome within a single live cell over time (Fig.4b, Fig.S10, Video 2), which score the spatial proximity according to the physical positions of genomic loci along the toroidal chromosome. The prominent domain boundary at the *dif* region was very persistent, and the weaker one at *oriC* region was present in the majority of the frames. Domain boundaries outside the *oriC*/*dif* regions were observed to change in distinct steps between consecutive frames. These proximity maps are reminiscent of Hi-C genomic-correlation maps, but present a significant advantage as they allow dynamic real-time mapping of a single genome in a live cell. The autocorrelation function of the density distributions along both the spatial (Fig.4c) and genomic coordinates (Fig.4d) decayed exponentially, with a decay half-time slightly smaller than 30 seconds, two orders of magnitude quicker than cell cycle time. Contrasting these fast density shifts, the full-image autocorrelation function of the chromosome morphology showed a much slower decay half-time, exceeding 5 minutes (Fig.4e). We thus find that whereas the global configuration of the *E. coli* chromosome is quite stable within a given cell boundary, local DNA arrangements are plastic, thus allowing, for example, fast regulations in gene expression to respond to environmental changes.

**Figure 4.**
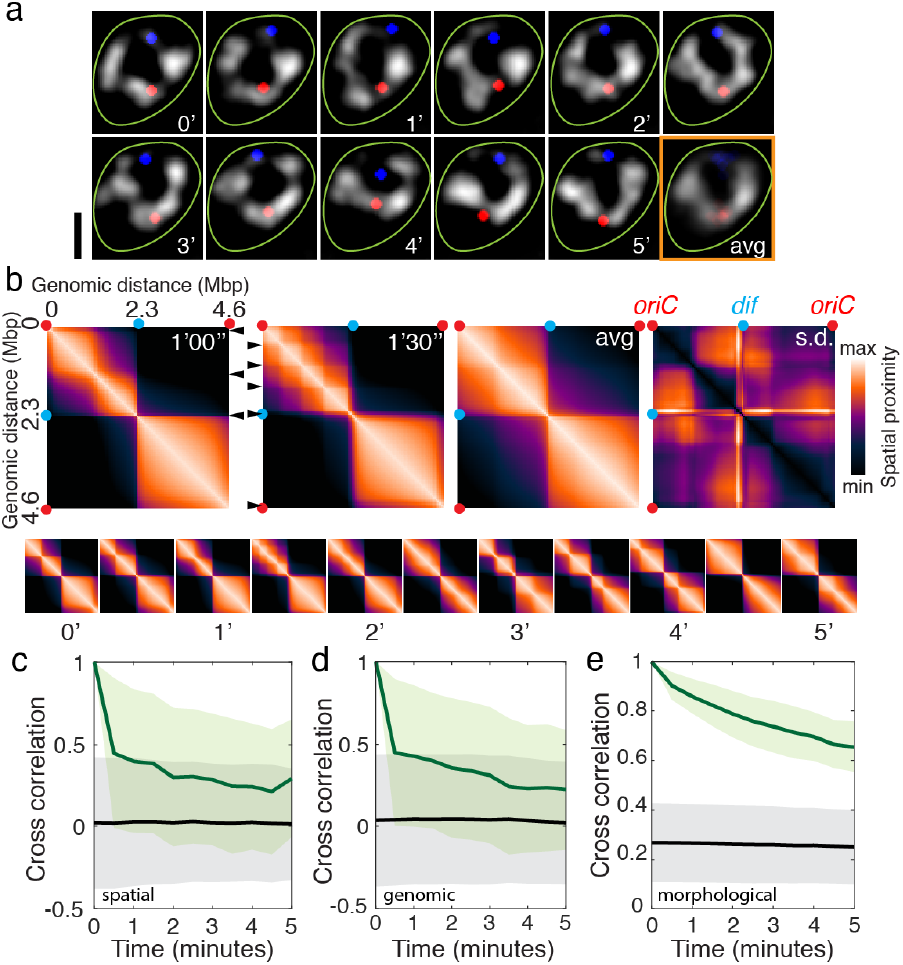
**Temporal dynamics of the circular chromosome of *E. coli*.** a. Time-lapse structured-illumination microscopy (SIM) images of a circular chromosome. Chromosomes in grey scale, Ori1/Ter3 localizations shown in red/blue, cell boundary in green. Time stamps are in minutes. The last frame is the time-average. Scale bar, 2 μm.
b. DNA spatial proximity maps derived from a single chromosome shown in a. Color bar indicates the level of spatial proximity for genomic loci along the circular chromosome. Arrow heads indicate apparent domain boundaries. Top panels, two consecutive frames, the time-average, and standard deviation values. Bottom panel, all 11 time frames. Red and cyan dots indicate Ori1/Ter3 loci.
c-e. Cross-correlation functions of individual chromosomes over time (green) compared to that across the cell population at each time point (black). These cross-correlation functions are measured with regard to the DNA density distribution along the contour in, respectively, spatial coordinates (c) genomic coordinates (d), and their morphology (e). Shown are mean and standard deviation values. n=46.

The direct imaging of the circular chromosome in live bacteria presented here provides a unique window to understanding the bacterial genome as a physical object. Specifically, we identified physical domain boundaries as key functional features. In contrast to all previous domain-based models where *oriC-* and *dif*-proximal regions were thought to form condensed domains^4,11,17,18^, we found persistent domain boundaries of reduced DNA density at the origin and terminus of replication. These *oriC-* and *dif*-proximal domain boundaries were found to be induced by MatP, which is involved in the spatiotemporal regulation of DNA segregation^25,27^, whereas other weak diffusive boundaries across the genome are governed by dynamic transcription processes. We expect that these physical-boundary-based principles may universally underlie the domain organization of bacterial chromosomes.

## Author contributions

FW and CD conceived and designed the project. FW constructed the bacterial strains. FW, AJ, and XZ did the microscopy experiments. JK led image analyses. JK, FW, and XZ wrote data analysis programs. All authors wrote the paper. CD supervised the project.

## Acknowledgements

We thank Jeremie Capoulade, Erwin van Rijn, Jelle van der Does, Louis Kuijpers, Margot Guurink, and Linda Chen (Huygens) for technical assistance, Jakub Wiktor for microscope upgrade and discussions, and David Sherratt, Rodrigo Reyes-Lamothe, and Jean-Luc Ferat for bacterial strains. This work was supported by ERC Advanced Grant SynDiv (No. 669598) and the Netherlands Organization of Scientific Research (NWO/OCW) as part of the Frontiers of Nanoscience Program. F.W. acknowledges support by Rubicon fellowship. A.J. acknowledges support by the Swiss National Science Foundation (Grants P2ELP2_168554 and P300P2_177768).

## Methods

**Strain construction.** For inducing a temperature sensitivity of DnaC, strain REP1329 (*dnaC2,ΔmdoB::Tn10*)^22^, a kind gift from Jean-Luc Ferat, was transformed with pKD46 and DNA fragment *ΔmdoB::aph frt* amplified from JW5794, to yield strain FW1957 (*dnaC2,ΔmdoB::aph frt*) through λ\RED recombination at 30°C. For endogenous HU labeling, strain FW1551 (*hupA-mYPet::aph frt)* was constructed and reported previously^31^. For fluorescence labeling of H-NS and Fis, linear fragments of *mYPet::aph frt* amplified from plasmid pROD61^32^ (a kind gift from the David Sherratt Lab), were transformed into strain W3110 to produce strain FW2561 (*hns-mYPet:: aph frt*) and strain FW2564 (*fis-mYPet::aph frt*), respectively, through λ\RED recombination^33^. For NAP deletions, strain JW3229-1 (*Δfis::aph frt*), JW1225-2 (*Δhns::aph frt*), JW0939 (*ΔmatP::aph frt*), JW3964(*ΔhupA::aph frt*), and JW0430(*ΔhupB::aph frt*) were obtained from the KeiO collection^34^. For experiments with Ori1/Ter3 labels, strain RRL189 (AB1157, *Ori1::lacOx240-hygR, Ter3::tetOx240-accC1ΔgalK::tetR-mCerulean frt, ΔleuB::lacI-mCherry frt*), a kind gift from Rodrigo Reyes-Lamothe, was used as a base to construct strains containing NAP labels and/or deletions through P1 phage transduction and cured of antibiotic resistance by flippase expressed from pCP20^33^. For L3/R3 labels, strain RRL66 (AB1157, *L3::lacOx240-hygR, R3::tetOx240-accC1 ΔgalK::tetR-mCerulean frt, ΔleuB::lacI-mCherry frt*), a kind gift from Rodrigo Reyes-Lamothe, was used as a base to construct strains like above. These strains were then finally transduced with the P1 phage produced from FW1957 to result in a DnaC2 temperature sensitivity. All strains used in this study are listed in Supplementary Table T1.

**Growth conditions.** For genetic engineering, *E. coli* cells were incubated in Lysogeny broth (LB) supplemented, when required, with 100 μg/ml ampicillin (Sigma-Aldrich), 50 μg/ml kanamycin (Sigma-Aldrich), or 34 μg/ml chloramphenicol (Sigma-Aldrich) for plasmid selection, and with 25 μg/ml kanamycin or 11 μg/ml chloramphenicol for selection of the genomic insertions of gene cassettes.

To obtain circular chromosomes, we grew cells in liquid M9 minimum medium (Fluka Analytical) supplemented with 2 mM MgSO_4_, 0.1mM CaCl_2_, 0.4% glycerol (Sigma-Aldrich), and 0.01% protein hydrolysate amicase (PHA) (Fluka Analytical) overnight at 30°C to reach late exponential phase. We then pipetted 1 μl culture onto a cover glass and immediately covered the cells with a flat agarose pad, containing the above composition of M9 medium as well as 6% agarose and 4 μg/ml A22, as described previously^35^. The cover glass was then immediately placed onto a baseplate and sealed with parafilm to prevent evaporation. The baseplate was placed onto the microscope inside a 40°C incubator for all cell growth and all imaging, unless noted differently. Circular chromosomes generally were imaged after 2.5-3 hours.

For treatment of circular chromosomes with antibiotics, the cells were inoculated in liquid M9 medium described above at 40°C with 4 μg/ml A22, and then placed under an agarose pad as described above with the addition of 100 μg/ml rifampicin or 10 μg/ml ciprofloxacin. The cells were incubated for 15 minutes before being imaged. Control samples did not have drugs added.

To reinitiate DNA replication, we grew the cells under agarose pad as described above for 3 hours, then moved the baseplate to room-temperature for 10 minutes before placing it back onto the microscope inside the 40°C chamber to prevent further new replication initiation and for imaging.

**Fluorescence imaging.** Wide-field Z scans were carried out using a Nikon Ti-E microscope with a 100X CFI Plan Apo Lamda Oil objective with an NA of 1.45. The microscope was enclosed by a custom-made chamber that was pre-heated overnight and kept at 40 °C. DAPI was excited by Nikon-Intensilight illumination lamp through a blue filter cube (λ_ex_ / λ_bs_ / λ _em_ =363-391 / 425 / 435-438 nm). mCerulean was excited by SpectraX LED (Lumencor) λ_ex_ = 430-450 through a CFP filter cube (λ_ex_ / λ_bs_ / λ_em_ =426-446 / 455 / 460-500 nm). mYPet signal was excited by SpectraX LED λ_ex_ = 510/25 nm through a triple bandpass filter λ_em_ = 465/25 – 545/30 – 630/60 nm. mCherry signals was excited by SpectraX LED λ_ex_ = 575/25 through the same triple bandpass filter. Fluorescent signals were captured by Andor Zyla USB3.0 CMOS Camera. In each channel, 19 slices were taken with a vertical step size of 227nm (in totally 4.3 μm). Structured Illumination Microscopy imaging was carried out using a Nikon Ti-E microscope and a SIM module. A 100X CFI Apo Oil objective with an NA of 1.49 was used. Samples were illuminated with 515nm laser line and a Nikon YFP SIM filter cube. mYPet, mCerulean, and mCherry signals of the same sample were also captured through wide-field imaging using a Nikon-Intensilight lamp. Filter cubes used for the wide-field imaging corresponding to the SIM images were CFP filters (λ_ex_ / λ_bs_ / λ _em_ =426-446 / 455 / 460-500 nm), YFP filters (λ_ex_ / λ_bs_ / λ_em =_ 490-510 / 515 / 520-550 nm), and RFP filters (λ_ex_ / λ_bs_ / λ_em_ = 540-580 / 585 / 592 - 668).

**3D deconvolution.** Image stacks of 19 slices were 3D-deconvolved using the Huygens Professional deconvolution software (Scientific Volume Imaging, Hilversum, The Netherlands), using an iterative Classic Maximum Likelihood Estimate (CMLE) algorithm with a point spread function (PSF) experimentally measured using 200nm multicolor Tetrabeads (Invitrogen). The PSF of the single-frame widefield images has a FWHM of 350 nm horizontally and 800 nm vertically. 3D deconvolution to a great extent reduced the out-of-focus noise in the images, which also lead to an improvement in lateral resolution. A deconvolved 200nm bead has FWHMs of 270 nm laterally and 580nm vertically. Due to the large vertical FWHM, inherent to wide-field imaging (including 3D-deconvolution), we find that the fluorescent signal at the central frame, rather than an integrated signal of all z frames, provides the best estimation of the local DNA density.

**Automated cell identification.** Phase contrast images were fed into a customized Matlab program to produce masks of cell boundaries, which then were used to allocate chromosomes and foci in other fluorescence channels. A manual correction and rejection process was carried out as a final step of quality control, to correct or reject cells when neighboring cells were too close to allow the automated program to distinguish. Chromosome foci numbers were then evaluated through methods as described previously^31^ to ensure that selected cells have a single chromosome copy.

**Automated blob analysis.** The blob analysis used an approach where a 3D-deconvolved image was subject to step-by-step stripping of the subsequent brightest Gaussian spots based on our measured PSF described above, until the image became blank (see Fig. S4a). The centers of the identified spots were then placed back into the image, where their mutual distances were evaluated (Fig. S4b). When the two spot centers were found to be located at a distance below our imaging resolution, they were assigned to the same blob (Fig. S4c). The average diameters and intensities of each blob were then measured for statistical analyses.

**Automated density analysis.** The center of mass of the circular chromosome was used to as the origin to create an angular coordinate system that assigns an angular (α) and radial (r) coordinate to each pixel. The circular chromosome was then sectioned along the angular axis into 100 bins, each 3.6° (Fig. S5a). The intensity maximum of each section was identified and mutually connected to constitute the ridge line of the filament. The points of this ridge line where then used to locate an improved center of mass using just these points. Using the new center, the ridge line was identified again. This process was iterated several times until the ridge line no longer was updated. Since the chromosome is not an isotropic torus, the identified points along the ridge line are not evenly spaced. In order to create evenly spaced coordinates, the filament was resampled with even spacing along the ridgeline. The total intensity of each section was then computed to represent the local intensity along the ridge line (where the ridge line was termed contour distance in the plots). To map the fluorescence signal at a particular position on the ridge line to the genomic sequence, we summed all intensity values and calculated the proportion of intensity that each section corresponded to (Fig. S5b-d). This proportion was thus translated into DNA content in each section and was used to map spatial coordinates into genomic coordinates as shown in Fig. 3d and Fig.S5d.

**Spatial proximity map construction.** An accumulative density function was constructed in a clockwise fashion along the ridge line of a single chromosome (Fig.S10a-d, also see Fig.3d), in which the contour distance between two genomic loci was estimated as indicated above. This spatial distance value was then plotted in the format of a color map on the pixels of the spatial proximity map (Fig.S10e).

All image analysis programs will be shared publicly on Github upon the publication of the paper.

